# Environmental, social, management and health factors associated with within- and between-individual variability in fecal glucocorticoid metabolite concentrations in zoo-housed Asian and African elephants

**DOI:** 10.1101/634691

**Authors:** Janine L. Brown, Jessica D. Bray, Kathy Carlstead, David Dickey, Charlotte Farin, Kimberly Ange-van Heugten

## Abstract

Identifying links between environmental, social, management, and health factors as they relate to physiological stress in captive elephants is crucial for the improvement of welfare and husbandry practices in North American zoos. Studies have examined the effects of short-term and chronic elevations in glucocorticoids in small groups of elephants, but few have examined adrenal activity on a large scale. This study evaluated 106 Asian *(Elephas maximus)* and 131 African *(Loxodonta africana)* elephants housed at 64 accredited facilities across North America. Fecal samples were collected every other week for 12 months and analyzed for glucocorticoid metabolite (FGM) concentrations. Risk factors for mean and individual variability (CV) in FGM were subjected to univariate and multivariable analyses using epidemiological methods. Independent variables that included individual traits, social environment, housing and management factors were chosen based on their identification as risk factors in previously published models for the same North American population of elephants. Results indicate that African elephants are more responsive to social stressors than Asians, and that poor joint health is a stress-related welfare problem for Asian, but not African elephants. For both species, higher FGM concentrations were associated with zoos located at more northern latitudes and having free access to indoor/outdoor spaces, whereas spending more time in managed interactions with staff were associated with lower FGM concentrations. Also important for captive management, elephants having diverse enrichment options and belonging to compatible social groups exhibited lower mean and reduced intra-individual variability in FGM. Our findings show that aspects of the zoo environment can be potential sources of stress for captive elephants, and that there are management activities that can facilitate coping and adapting to zoo conditions. Given species differences in factors that affected FGM, targeted, species-specific management approaches likely are needed to ensure good welfare for all elephants.

## Introduction

Modern zoos strive to ensure animals under human care experience a high standard of welfare that meets emotional and physical health needs [1]. Asian (*Elephas maximus*) and African (*Loxodonta africana*) elephants in zoos have received considerable scrutiny in the last decade due to concern over welfare and management practices [2]. To be successful, it is important that captive elephant programs evaluate the basic husbandry needs of individual animals, as well as the more complex factors that may affect welfare in a captive environment. An earlier study found no differences in serum cortisol concentrations or cortisol variability in elephants managed in either free contact (elephants and people share the same space) or protected contact (elephants and people are separated by a barrier); however, there was a significant facility effect [3], suggesting that facility-specific differences in management exist that may affect adrenal activity and cortisol levels in captive elephants.

A recent ‘Elephant Welfare Project (EWP)’ took an epidemiological approach to investigating the factors that impact zoo elephant welfare in North America [4]. That study, conducted by a multi-institutional team of researchers and including 255 elephants at 68 Association of Zoos and Aquariums (AZA) accredited zoos, found that herd social structure, caretaker interactions, and enrichment, and feeding diversity correlated with a variety of welfare outcomes [5–16]. In particular, enrichment and social factors were important for reproductive activity and reducing stereotypic behaviors, diversity of feeding practices and exercise reduced the likelihood that an elephant would be overweight, softer exhibit substrates were good for physical and behavioral health, and positive keeper-elephant relationships were mutually beneficial. Overall, environments that provided diversity and choice were of greater importance to elephant welfare than exhibit size alone. A remaining question is how these factors affect physiological stress responses in individual elephants, and their ability to cope with a zoo environment.

The most commonly used bio-markers of stress and, by extension welfare, are glucocorticoids (GC) that are secreted from the adrenal gland in response to a stressor [17]. Both favorable and aversive stimuli can stimulate GC release; eustress defines responses beneficial to an animal’s wellbeing [19], while distress indicates a negative reaction to a stressor [18]. To add to the physical [12,13], behavioral [7–9], and physiological [5,15] outcomes measured in the EWP to date, assessing how factors in the zoo environment affect GC responses would benefit from a similar epidemiological approach. Prolonged exposure to psychological or physical stressors, and chronic elevations in GCs, can result in immunosuppression, decreased wound healing, increased susceptibility to disease, poor reproduction, and development of stereotypic behaviors [17]. Glucocorticoid concentrations in blood samples are one indicator of adrenal activity in response to a stressor [21,21] and have been measured in wild and captive elephants [15,22,23,24]. However, there are limitations to using blood GCs as an index of stress if the act of collecting the sample itself elicits a response [25,26]. Development of noninvasive techniques to measure GCs or their metabolites excreted in feces (FGM) has provided us with a robust tool for wildlife studies, including in elephants [27–31]. Non-invasive FGM monitoring has been applied to studies of welfare across a diverse array of species [33,34], including elephants [35,36], and aided in improving *ex situ* management [37–39].

The goal of this study was to use multi-variable modeling to assess if the already-identified management, facility, keeper, enrichment, individual, or social factors that are associated with other welfare outcomes for elephants [5,7–9,12,13,15] also are risk factors for elevated FGM concentrations. Recently, Edwards et al. [16] found positive correlations between the number of clinical cases in the 1-year EWP study and the coefficient of variation (CV) for both serum cortisol and FGM, suggesting that within-individual variation in FGMs can be a welfare indicator of stress-related pathology. The goal of this study was to better understand relationships between FGM and welfare outcomes, and how they are influenced by extrinsic forces – important information needed to optimize management of elephants in zoo settings.

## Materials and methods

### Ethics statement

This research was approved by the management at each participating institution, and where applicable, was reviewed and approved by zoo research committees. In addition, the study protocol was approved by the Smithsonian National Zoo (NZP-ACUC #11/10).

### Study population and sample collection

The study consisted of 237 captive elephants, 106 Asian (85 females; 21 males) and 131 African (104 females; 27 males), housed at 64 American Zoo and Aquarium (AZA) accredited facilities throughout North America that participated in the EWP [4]. Fecal samples were collected every other week for 12 months. The sampling protocol required samples to be collected fresh from the ground, mixed to obtain homogeneity, and then 5-10 subaliquots (~50-100 g) placed into Whirlpak^®^ plastic bags, and frozen (−20°C) immediately.

### Fecal extraction and GC metabolite analysis

Fecal samples were lyophilized (Labconco, Kansas City, MO), and 0.1 g (+/− 0.02) of well-mixed fecal powder was placed into 16 x 125 mm glass tubes (Fisher Scientific; Pittsburgh, PA). Five ml of 80% methanol was then added and the samples were mixed for 30 minutes on a multi-tube vortexer (Glas-Col; Terre Haute, IN), followed by centrifugation for 20 min at 2500 x g (Sorvall RC 3C Plus; Thermo Fisher Scientific, Waltham, MA). Each supernatant was recovered and the remaining pellet was resuspended in 5 ml of 80% methanol and extracted again. The two supernatants were combined into a 16 x 125 mm glass tubes and dried under forced air in a fume hood overnight. Extracted samples were reconstituted in 1 ml of 100% methanol, dried again, and then buffer (1 ml, 0.149 M NaCl, 0.1 M NaPO_4_; with pH 7.0) added and the tubes sonicated (Part# 08895-60; Cole-Parmer, Vernon Hills, IL) for 30 seconds to dissolve particulates. Finally, all samples were diluted (1:8) in assay buffer (Cat. No. X065, Arbor Assays, Arbor, MI, USA) and stored at –20°C until enzyme immunoassay (EIA) analysis.

Concentrations of FGM were determined using a double-antibody enzyme EIA with a polyclonal rabbit anti-corticosterone antibody (CJM006) validated for elephants [40]. Standards (3.9-1000 pg/well; Sigma Diagnostics, St. Louis, MO), samples, and controls were added in duplicate (50 μl per well) to pre-coated goat anti-rabbit IgG, 96-well plates at room temperature. Corticosterone-horseradish peroxidase (25 μl, 1:20,000 dilution) was immediately added to all wells, followed by 25 μl anti-corticosterone antibody (1:60,000) that was added to all but non-specific binding wells. The plates were covered with microplate sealers and incubated at room temperature on an agitator (Model E6121; Eberbach Corp., Ann Arbor, MA) for 1 hour. All plates were then washed four times (1:20 dilution, 20X Wash Buffer Cat. No. X007; Arbor Assays), blotted dry, and 100 μl of TMB (3, 3’, 5, 5’ – tetramethylbenzidine) (Moss Inc., Pasadena, MD) was added. Plates were incubated for 30-45 min at room temperature without shaking, and the reaction stopped by adding 50 μL of a 1 N HCl solution. Optical density was read in a plate reader at 450 nm (OPsys MR; Dynex Technologies, Chantilly, VA). The inter-assay coefficient of variation (CV %) for the high control was 8.1%, and the low control CV% was 15.1% (n=200 plates); intra-assay CV was <10% as all samples with duplicate CVs over 10% were reanalyzed. Assay sensitivity (based on 90% binding) was 0.14 ng/ml.

### Statistical Analysis

#### Independent Variables

Independent variables used for these analyses were chosen based on their significance in already-published multi-variable models for other “gold standard” welfare indicators of the EWP (ovarian cyclicity, stereotypy, body condition, foot and joint health, walking distance and recumbency, and serum cortisol). Full details regarding data collection and variable creation are provided in earlier publications [5–16]. Table 1 lists the welfare indicators and descriptions of the independent variables. For ease of discussion, independent variables were categorized as measures of Individual traits, Social environment, Housing factors or Management variables. There were two levels of measurements for independent variables: individual elephant and zoo-level. Elephant-specific independent variables were: *Age, Sex, Percent Time in Mixed-Sex Herds, Social Group Contact, Walking Hours Per Week, Percent Time with Juveniles, Percent Time Housed Separately, Transfers, Percent Time In/Out Choice, Social Experience, Recumbence Rate, Percent Time on Hard Substrate, Percent Time on Soft Substrate, Space Experience Outdoors at Night, Space Experience with In/Out Choice, Joint Health, Space Experience Total at Night, Mean Daily Walking Distance, Mean Serum Cortisol, Elephant Positive Behaviors,* and *Elephant Interacts with Public.* Measured on a zoo-level were *Season*, *Enrichment Diversity, Alternative Feeding Methods, Feeding Diversity, Percent Time Managed, Keeper Positive Opinions of Elephants, Keeper as Herdmate* and *Latitude of Zoo*.

**Table 1.** Variable significant independent variables, for either or both species, in multi-variable models of welfare outcomes from the Elephant Welfare Project. Groups: S=social, H=housing, M=management, I=individual.

| Welfare Indicators | Independent Variable | Group | Definition |
| --- | --- | --- | --- |
| Ovarian Cycling<sup>1</sup> | Percent Time in Mixed Sex Herds (*unpub.*) | S | Sum of monthly percent time spent in social groups where both males and females are present |
| Prolactin<sup>1</sup> | Enrichment Diversity | M | Shannon diversity index score of enrichment activities types and frequencies conducted at zoo |
|  | Alternate Feeding Methods | M | The proportion of all feedings where food was presented in a foraging device, hidden, or hung above the exhibit |
|  | Social Group Contact | S | Maximum number of unique social groups focal animal is part of |
| Body Condition<sup>2</sup> | Walking, Hours/Week | M | Number of reported hours spent walking elephants each week, ranging from 1 (< 1 hour per week) to 7 (14 or more hours per week) |
|  | Feeding Diversity | M | Shannon diversity index score of feeding types and frequencies conducted at zoo |
|  | Sex (ref: male) | I | Male or female |
| Daytime Stereotypy<sup>3</sup> | Percent Time Managed | M | Sum of percent time spent in activities managed by caretaking staff |
|  | Percent Time with Juveniles | S | Sum of monthly percent time spent in social groups where an elephant 7 years or younger was present |
|  | Percent Time Housed Separately | S | Sum of monthly percent time spent housed in a social group of one |
|  | Transfers | I | Total number of inter-zoo transfers an elephant has experienced |
| Nighttime Stereotypy<sup>3</sup> | Percent Time In/Out Choice | M | Sum of monthly percent time spent in environments where there is a choice of indoors or outdoors |
|  | Social Experience | S | The average weighted (by percent time) size of all social groups in which an elephant spent time |
| Recumbence<sup>4</sup> | Recumbence Rate | I | Hours recumbent per day, averaged over all days of data collection |
|  | Percent Time on Hard Substrate | H | Sum of monthly percent time spent in environment with 100% concrete or stone aggregate substrate |
|  | Percent Time Soft Substrate | H | Sum of monthly percent time spent in environment with 100% grass, sand, or rubber substrate |
|  | Space Experience Outdoor Night (per 500 ft<sup>2</sup>) | H | The average weighted (by percent time) size of all environments in which an elephant spent time in outdoor environments only |
|  | Percent Time Housed Separately | H | Sum of monthly percent time spent housed in a social group of one |
| Muscoskeletal Score<sup>5</sup> | Space Experience In/Out Choice (per 500ft<sup>2</sup>) | S | The average weighted (by percent time) size of all environments in which an elephant spent time where there is a choice of indoors or outdoors |
|  | Joint Abnormalities (ref: absence) | I | Presence or absence of gait change, limb deformity, joint heat or swelling noted from muscoskeletal exam |
| Foot Health<sup>5</sup> | Percent Time In/Out Choice | M | Sum of monthly percent time spent housed in a social group of one |
|  | Space Experience Total Night (per 500 ft<sup>2</sup>) | H | The average weighted (by percent time) size of all environments in which an elephant spent time at night |
|  | Age | I | Age of elephant in years in 2012 |
| Walking Distance<sup>6</sup> | Mean Daily Walking Distance | I | Mean outdoor daily walking distance measured by anklets equipped with GPS data loggers |
|  | Social Group Contact | S | Maximum number of unique social groups focal animal is part of |
|  | Feeding Predictability (ref: unpredictable) | M | The predictability of feeding times; categorical where 1 is predictable, 2 is semi-predictable, and 3 is unpredictable |
|  | Space Experience Total Night (per 500 ft<sup>2</sup>) | H | The average weighted (by percent time) size of all environments in which an elephant spent time in outdoor environments only |
| Serum Cortisol<sup>7</sup> | Mean Serum Cortisol | I | Mean of 24 blood samples taken bi-weekly for 1 year |
|  | Keeper Attitude: Positive Opinions of Elephants | M | Composite scores (averaged by zoo) of keepers' opinions of elephants: elephants are playful, like to be trained, like change, are trusting, affectionate, and bond to keepers |
|  | Keeper Attitude: Keeper as Herdmate | M | Composite scores (averaged by zoo) of keepers' perceptions that they are accepted by elephants as part of the herd, elephants are interested in the keepers, keepers connect verbally with elephants, keepers have bonds with elephants |
|  | Latitude of Zoo | H | Angular distance of a zoo's location north of the equator |
|  | Elephant Positive Behaviors | I | Composite scores (from keeper ratings) for affiliative/friendly behaviors, food sharing, solo play, wallowing |
|  | Elephant Interacts with Public | I | Composite scores (from keeper ratings) for elephant initiates watches and initiates interactions with zoo visitors |

<sup>1</sup>Brown et al. [5]; <sup>2</sup>Morfeld et al. [13]; <sup>3</sup>Greco et al. [7]; <sup>4</sup>Holgate et al. [9]; <sup>5</sup>Miller et al. [12]; <sup>6</sup>Holgate et al. [8]; <sup>7</sup>Carlstead et al. [15].

Generalized Linear Mixed Models (GLMM) were used to determine *Species* and *Season* effects on mean FGMs, and *Species* and *Sex* effects on mean and CV of FGMs; Zoo was treated as a random effect to account for clustering of elephants by facility. Mean FGM concentrations for elephants of each species, and CV of FGMs for both species combined, were fitted in regression models using Generalized Estimating Equations (GEE), which allow for the individual elephant to be used as the unit of analysis, accounts for clustering of individuals within zoos, and focuses on population-averaged effects [41]. The model included repeated measures of FGMs by *Season.* Zoos were treated as random effects and an independent correlation structure was specified. We built multi-variable regression models by first assessing individual predictors at the univariate level and then at the bivariate level with each demographic variable (*Species, Age, Sex)* as potential confounding variables. Confounding variables (those that altered the beta values of input variables by more than 10% during bivariate analysis) were included in all models as necessary. Any variables that predicted FGM mean or CV (P < 0. 15) following the univariate and bivariate assessments were retained for evaluation in the hierarchical model building process. The hierarchical selection was based on quasi-likelihood under the independence model criterion (QIC) values and parameter estimates of explanatory variables. Models exhibiting multi-collinearity, as defined by a variance inflation factor of greater than 10 and a Condition Index of greater than 30, were not considered for further analysis.

Unless otherwise indicated, differences were considered significant at P < 0.05. All analyses were conducted using IBM SPSS Statistics Version 25, IBM Corp., Armonk, NY, USA.

## Results

The elephant study population ranged in age from 0 to 64 years (mean age: Asian, 34.3 ±1.5 years; African, 27.7 ±1.1). Table 2 presents seasonal mean FGM concentrations for each species. Overall FGM concentrations were higher in Asian (124.41 ± 4.89 ng/g) than African (97.73 ± 3.01 ng/g) elephants. There was a significant main effect of species (F = 27.86, P = 0.000), but not season (F = 1.30, P = 0.000). In all seasons, Asian elephants had higher mean concentrations than Africans.

**Table 2.** Mean (± SEM) and minimum-maximum seasonal fecal glucocorticoid metabolite concentrations in Asian (n = 106) and African (n = 131) elephants in North American zoos that participated in the Elephant Welfare Project.

| Season | Asian Elephants |  |  | African Elephants |  |  |
| --- | --- | --- | --- | --- | --- | --- |
| | Mean ± SEM | Min | Max | Mean ± SEM | Min | Max |
| Winter (Jan-Mar) | 146.9 ± 5.01 | 43.41 | 317.67 | 108.48 ± 3.03 | 31.83 | 222.49 |
| Spring (Apr-Jun) | 156.8 ± 5.04 | 57.78 | 286.74 | 107.22 ± 3.01 | 37.56 | 266.17 |
| Summer (Jul-Sep) | 146.2 ± 4.27 | 49.74 | 324.18 | 105.04 ± 2.94 | 28.81 | 229.71 |
| Fall (Oct-Dec) | 147.8 ± 5.13 | 37.82 | 310.56 | 110.01 ± 3.08 | 26.78 | 292.43 |

Mean and average variability (CV) of FGMs was calculated for the entire year and is given for each species and sex separately in Table 3. GLMM analysis demonstrates significant differences for *Species* (F=8.496, P=0.004), but not for *Sex* (F=0.124, P=0.726, Table 3). For FGM CV, which is a normalized calculation, there were no significant effects of *Species* (F=0.004, P=0.950) or *Sex* (f=0.891, P=0.346). Therefore, mean FGMs were analyzed separately for each species, whereas FGM CVs were analyzed for both species combined. Descriptive statistics for independent variables are presented for each species in Table 4.

**Table 3.** Mean (± SEM) fecal glucocorticoid metabolite (FGM) concentrations and coefficient of variation (CV) for male and female Asian and African elephants in North American zoos that participated in the Elephant Welfare Project.

|  | Asian Elephants |  | African Elephants |  |
| --- | --- | --- | --- | --- |
|  | Male = 21 | Female = 85 | Male = 27 | Female = 104 |
| Mean FGM (ng/g) | 121.55 ± 8.69 | 125.47 ± 4.87 | 99.6 ± 5.70 | 97.7 ± 3.14 |
| Mean FGM CV | 31.53 ± 1.49 | 32.44 ± 1.28 | 35.20 ± 2.55 | 33.17 ± 1.18 |

**Table 4.** Descriptive statistics (mean, SEM, minimum, maximum) for independent variables of Asian and African elephants in North American zoos that participated in the Elephant Welfare Project.

|  | Asian Elephants |  |  |  |  |  | African Elephants |  |  |  |  |
| --- | --- | --- | --- | --- | --- | --- | --- | --- | --- | --- | --- |
|  | N | Mean | SEM | Min | Max |  | N | Mean | SEM | Min | Max |
| Fecal Glucocorticoid Metabolites - Mean | 106 | 124.69 | 4.26 | 59.69 | 282.88 |  | 131 | 98.11 | 2.75 | 40.56 | 211.34 |
| Fecal Glucocorticoid Metabolites - CV | 106 | 32.26 | 1.07 | 9.78 | 71.24 |  | 131 | 33.59 | 1.070 | 15.20 | 92.59 |
| Percent Time in Mixed Sex Herds | 106 | 12.46 | 2.969 | 0.00 | 100.00 |  | 131 | 23.31 | 3.200 | 0.00 | 100.00 |
| Enrichment Diversity | 93 | 2.91 | 0.015 | 2.54 | 3.16 |  | 129 | 2.83 | 0.014 | 2.54 | 3.26 |
| Alternate Feeding Methods | 100 | 0.49 | 0.022 | 0.08 | 0.92 |  | 131 | 0.38 | 0.019 | 0.08 | 0.91 |
| Social Group Contact | 106 | 2.70 | 0.200 | 1.00 | 11.00 |  | 131 | 4.94 | 0.618 | 1.00 | 30.00 |
| Walking, Hours/Week | 88 | 2.58 | 0.186 | 1.00 | 7.00 |  | 129 | 1.92 | 0.130 | 1.00 | 7.00 |
| Feeding Diversity | 95 | 1.37 | 0.032 | 0.31 | 1.78 |  | 129 | 1.38 | 0.018 | 0.98 | 1.79 |
| Sex (ref: male) | 106 | 0.80 | 0.039 | 0.00 | 1.00 |  | 131 | 0.79 | 0.035 | 0.00 | 1.00 |
| Percent Time Managed | 89 | 55.42 | 2.035 | 20.00 | 91.00 |  | 129 | 49.34 | 1.640 | 13.00 | 100.00 |
| Percent Time with Juveniles | 106 | 18.63 | 3.413 | 0.00 | 100.00 |  | 131 | 22.78 | 3.310 | 0.00 | 100.00 |
| Percent Time Housed Separately | 106 | 32.96 | 3.817 | 0.00 | 100.00 |  | 131 | 21.15 | 2.590 | 0.00 | 100.00 |
| Transfers | 106 | 2.69 | 0.204 | 0.00 | 10.00 |  | 129 | 2.68 | 0.162 | 0.00 | 10.00 |
| Percent Time In/Out Choice | 106 | 15.74 | 2.157 | 0.00 | 77.67 |  | 131 | 17.30 | 1.820 | 0.00 | 89.82 |
| Social Experience | 106 | 2.17 | 0.106 | 1.00 | 4.93 |  | 131 | 3.14 | 0.218 | 1.00 | 11.22 |
| Recumbence Rate | 25 | 8.02 | 0.752 | 0.00 | 19.72 |  | 38 | 5.34 | 0.452 | 0.05 | 9.17 |
| Percent Time on Hard Substrate | 106 | 9.69 | 1.260 | 0.00 | 51.80 |  | 131 | 13.13 | 1.080 | 0.00 | 50.00 |
| Percent Time Soft Substrate | 106 | 10.82 | 1.228 | 0.00 | 55.90 |  | 131 | 10.61 | 1.260 | 0.00 | 58.30 |
| Space Experience Outdoor Night (per 500 ft<sup>2</sup>) | 106 | 34.60 | 3.903 | 0.00 | 187.39 |  | 131 | 70.75 | 8.910 | 0.00 | 574.28 |
| Space Experience In/Out Choice (per 500 ft<sup>2</sup>) | 106 | 19.36 | 2.177 | 0.00 | 92.13 |  | 131 | 38.35 | 5.560 | 0.00 | 312.74 |
| Joint Abnormalities (ref: absence) | 98 | 0.33 | 0.048 | 0.00 | 1.00 |  | 94 | 0.23 | 0.044 | 0.00 | 1.00 |
| Space Experience Total Night (per 500 ft<sup>2</sup>) | 106 | 27.64 | 2.760 | 1.09 | 147.05 |  | 131 | 56.25 | 6.920 | 0.88 | 419.14 |
| Age of Elephant | 106 | 34.84 | 1.459 | 1.00 | 64.00 |  | 131 | 27.85 | 1.060 | 0.00 | 52.00 |
| Mean Daily Walking Distance | 26 | 5.31 | 0.629 | 1.21 | 17.26 |  | 34 | 5.42 | 0.260 | 2.19 | 9.71 |
| Feeding Predictability (ref: unpredictable) | 95 | 2.16 | 0.066 | 1.00 | 3.00 |  | 129 | 1.93 | 0.050 | 1.00 | 3.00 |
| Mean Serum Cortisol | 98 | 17.83 | 0.748 | 5.96 | 40.02 |  | 115 | 17.95 | 0.583 | 5.87 | 37.26 |
| Keeper Attitude: Positive Opinions of Elephants | 84 | 3.68 | 0.053 | 1.59 | 4.40 |  | 106 | 3.65 | 0.050 | 2.77 | 5.37 |
| Keeper Attitude: Keeper as Herdmate | 84 | 3.02 | 0.073 | 2.00 | 4.48 |  | 106 | 2.65 | 0.054 | 1.41 | 4.03 |
| Latitude of Zoo | 103 | 35.81 | 0.567 | 21.00 | 47.00 |  | 131 | 35.60 | 0.414 | 26.00 | 47.00 |
| Elephant Positive Behaviors | 67 | 4.45 | 0.128 | 1.53 | 6.31 |  | 93 | 4.67 | 0.080 | 2.21 | 6.42 |
| Elephant Interacts with Public | 67 | 2.48 | 0.107 | 0.98 | 5.68 |  | 93 | 2.40 | 0.082 | 0.83 | 5.16 |

For Asian and African elephants separately, univariate linear regressions of independent variables with mean FGM concentrations are shown in Table 5. For Asians, significant negative associations (i.e., lower FGMs) were observed for *Enrichment Diversity, Walking (hrs/week), Percent Time Managed by Staff, Experience Outdoors at Night, Space Experience with In/Out Choice, Total Space Experienced at Night, Mean Daily Walking Distance* and *Latitude of Zoo.* Positive associations (i.e., higher FGMs) were associated with *Percent Time Housed Separately, Recumbent Rate, Joint Abnormalities, Serum Cortisol* and *Keeper as Herdmate.* For Africans, significant negative regressions with mean FGMs were with *Percent Time Managed* (as with Asians), and *Percent Time with In/Out Choice,* and additionally with *Keeper as Herdmate.* Positive associations were with *Percent Time in Mixed Sex Herds, Social Experience, Social Group Contact, Feeding Predictability, Latitude of Zoo, Mean Daily Walking Distance,* and all three *Space Experience* variables. Overall, African FGMs were associated with three social variables and only one individual variable (*Mean Daily Walking Distance),* whereas FGMs in Asians were associated with only one social variable (*Percent Time Housed Separately)* and four individual variables. Both species had equal numbers of management and housing variables associated with FGM concentrations.

**Table 5.** Univariate linear regressions of 12-month mean fecal glucocorticoid metabolite concentrations in Asian and African elephants in North American zoos and previously published risk factors (independent variables) from the Elephant Welfare Project. Variables entered into multi-variable analyses (P<0.15) are **bolded**.

|  | Asian Elephants |  |  |  | African Elephants |  |  |  |
| --- | --- | --- | --- | --- | --- | --- | --- | --- |
| **Independent Variable** | **N** | **Estimate** | **SEM** | **P value** | **N** | **Estimate** | **SEM** | **P value** |
| Percent Time in Mixed Sex Herds (*unpub.*) | 106 | -0.065 | 0.140 | 0.646 | **131** | **0.211** | **0.073** | **0.005** |
| Enrichment Diversity | **93** | **-58.746** | **31.058** | **0.062** | 129 | 14.139 | 16.989 | 0.407 |
| Alternate Feeding Methods | 100 | 16.049 | 20.348 | 0.432 | 131 | 13.994 | 12.529 | 0.266 |
| Social Group Contact | 106 | -0.312 | 2.088 | 0.882 | **131** | **0.944** | **0.383** | **0.015** |
| Walking, Hours/Week | **88** | **-4.796** | **2.673** | **0.076** | 129 | -2.274 | 1.864 | 0.225 |
| Feeding Diversity | 95 | -10.397 | 14.750 | 0.483 | 129 | 8.369 | 13.265 | 0.529 |
| Sex (ref: male) | 106 | 3.971 | 10.721 | 0.712 | 133 | -1.543 | 6.804 | 0.821 |
| Percent Time Managed | **89** | **-0.545** | **0.253** | **0.034** | **128** | **-0.284** | **0.149** | **0.060** |
| Percent Time with Juveniles | 106 | -0.043 | 0.122 | 0.726 | 131 | 0.079 | 0.073 | 0.283 |
| Percent Time Housed Separately | **106** | **0.174** | **0.108** | **0.109** | 131 | 0.023 | 0.093 | 0.804 |
| Transfers | 106 | -0.964 | 2.040 | 0.637 | 131 | -0.852 | 1.479 | 0.566 |
| Percent Time In/Out Choice | 106 | -0.074 | 0.188 | 0.695 | **131** | **-0.285** | **0.166** | **0.088** |
| Social Experience | 106 | -5.197 | 3.918 | 0.188 | **131** | **2.342** | **1.089** | **0.033** |
| Recumbence Rate | **25** | **4.949** | **2.200** | **0.034** | 38 | 0.908 | 1.639 | 0.583 |
| Percent Time on Hard Substrate | 106 | 0.725 | 0.323 | 0.027 | 131 | 0.132 | 0.223 | 0.556 |
| Percent Time Soft Substrate | 106 | -0.115 | 0.340 | 0.735 | 131 | 0.229 | 0.190 | 0.229 |
| Space Experience Outdoor Night (per 500 ft<sup>2</sup>) | **106** | **-0.187** | **0.105** | **0.080** | **131** | **0.073** | **0.026** | **0.006** |
| Space Experience In/Out Choice (per 500 ft<sup>2</sup>) | **106** | **-0.333** | **0.189** | **0.081** | **131** | **0.110** | **0.042** | **0.010** |
| Joint Abnormalities (ref: absence) | **95** | **20.198** | **7.470** | **0.008** | 96 | 0.298 | 7.660 | 0.969 |
| Space Experience Total Night (per 500 ft<sup>2</sup>) | **106** | **-0.282** | **0.149** | **0.060** | **131** | **0.111** | **0.033** | **0.001** |
| Age of Elephant | 106 | 0.261 | 0.285 | 0.361 | 133 | -0.278 | 0.227 | 0.222 |
| Mean Daily Walking Distance | **26** | **-5.144** | **2.380** | **0.041** | **34** | **6.428** | **3.264** | **0.058** |
| Feeding Predictability (ref: unpredictable) | 95 | 0.642 | 7.087 | 0.928 | **129** | **6.221** | **4.167** | **0.138** |
| Mean Serum Cortisol | **98** | **1.208** | **0.591** | **0.024** | 117 | 0.196 | 0.475 | 0.680 |
| Keeper Attitude: Positive Opinions of Elephants | 84 | 6.814 | 10.654 | 0.524 | 108 | -3.814 | 4.838 | 0.432 |
| Keeper Attitude: Keeper as Herdmate | **84** | **16.663** | **7.625** | **0.032** | **108** | **-10.227** | **4.683** | **0.031** |
| Latitude of Zoo | **106** | **-1.153** | **0.665** | **0.086** | **133** | **1.659** | **0.563** | **0.004** |
| Elephant Positive Behaviors | 67 | -5.672 | 4.667 | 0.229 | 93 | -0.505 | 3.728 | 0.893 |
| Elephant Interacts with Public | 67 | -0.212 | 5.644 | 0.970 | 93 | 0.503 | 3.639 | 0.890 |

Multivariable analyses required the exclusion of *Mean Daily Walking Distance* and *Recumbent Rate* because these variables were measured in only a sub-set of the elephants. Also, *Social Experience* was highly correlated (r = 0.899) with *Social Group Contact* and so also was not included in the multivariable model building process due to collinearity problems. The final models are given in Table 6 for Asian and Table 7 for African elephants.

**Table 6.** Multi-variable model of seasonal fecal glucocorticoid metabolite concentrations for Asian elephants (n=106) in North American zoos that participated in the Elephant Welfare Project^1^.

| Variable | Beta Estimate | SEM | P value |
| --- | --- | --- | --- |
| Intercept | 118.69 | 23.60 | 0.000 |
| *Season*: Winter (Jan-Mar) | -2.43 | 24.92 | 0.922 |
| *Season*: Spring (Apr-Jun) | -42.59 | 24.01 | 0.076 |
| *Season*: Summer (Jul-Sep) | -10.91 | 21.21 | 0.606 |
| *Season*: Fall (Oct-Dec) (ref) | 0 |  |  |
| Sex: Female | -3.15 | 6.83 | 0.644 |
| Sex: Male (ref) | 0 |  |  |
| Age of Elephant | 0.34 | 0.22 | 0.128 |
| Joint Health: no abnormalities | -21.14 | 8.58 | 0.014 |
| Joint Health: abnormalities (ref) | 0 |  |  |
| Space Experience In/Out Choice (per 500 ft<sup>2</sup>) | -0.41 | 0.13 | 0.003 |
| *Season*: Winter*Latitude of Zoo | 0.61 | 0.66 | 0.350 |
| *Season*: Spring* Latitude of Zoo | 1.81 | 0.77 | 0.019 |
| *Season*: Summer*Latitude of Zoo | 0.66 | 0.62 | 0.288 |
| *Season*: Fall*Latitude of Zoo (ref) | 0.39 | 0.55 | 0.473 |

<sup>1</sup>Age is a confounder for *Sex* and *Latitude of Zoo*.

**Table 7.** Multi-variable model of seasonal fecal glucocorticoid metabolite concentrations for African elephants (n=131) in North American zoos that participated in the Elephant Welfare Project^1^.

|  | **Beta Estimate** | **SEM** | **P value** |
| --- | --- | --- | --- |
| Intercept | 16.67 | 26.24 | 0.525 |
| *Season: Winter (Jan-Mar)* | -3.79 | 2.94 | 0.197 |
| *Season: Spring (Apr-Jun)* | -1.10 | 3.03 | 0.716 |
| *Season: Summer (Jul-Sep)* | -1.71 | 2.80 | 0.541 |
| *Season: Fall (Oct-Dec) (ref)* | 0 |  |  |
| Sex: Female | -5.53 | 6.69 | 0.409 |
| Sex: Male (ref) | 0 |  |  |
| Age | -0.10 | 0.28 | 0.719 |
| Percent Time Managed | -0.27 | 0.13 | 0.045 |
| Latitude of Zoo | 2.62 | 0.58 | 0.000 |
| Percent Time in Mixed-Sex Herds | 0.19 | 0.09 | 0.039 |
| Space Experience Outside at Night (per 500 ft<sup>2</sup>) | 0.06 | 0.02 | 0.004 |
| Percent Time In/Out choice | -0.20 | 0.09 | 0.032 |

<sup>1</sup>*Age of elephant is a confounder of Percent Time Managed and Latitude of Zoo. Latitude of Zoo was a confounder of Percent Time in Mixed-Sex Herds and Space Experience Outside at Night.*

The multi-variable model for Asian elephant FGMs included both Season and *Latitude of Zoo* with almost significant main effects (P = 0.076 and 0.051, respectively), so they were also added as an interaction term, *Season*Latitude of Zoo*, in the model (Table 6). The rationale for this was that the degree of climatological change between seasons is a function of how far north the zoo lies. This interaction factor was a significant risk factor for higher FGM: spring season at higher latitudes. When all other independent variables are held constant, an increase of one degree in *Latitude of Zoo* corresponds to a 1.81 ng/g increase in FGM during April – June. For Asian elephants, risk factors for higher FGMs were *Joint Abnormalities* and limited *Space Experience with In/Out Choice*. Our analysis found that, when all other independent variables are held constant, the absence of *Joint Abnormalities* decreases FGM by 21.14 ng/g, and for every 5000 ft^2^ increase in *Space Experience with In/Out Choice* there is a 4.10 ng/g decrease in FGM.

The multivariable model for African elephant FGMs also demonstrated effects of *Latitude of Zoo* on FGM, but no seasonal effects (Table 6). As latitude increases by one degree, FGMs increase by 2.67 ng/g. There were four additional risk factors in the multivariable model: *Percent Time In/Out Choice*, and *Percent Time Managed* by staff. For every 10% increase in *Percent Time In/Out Choice* there is a 2.00 ng/g decrease in FGM. Similarly, a 10% increase *Percent Time Managed* decreases FGMs by 2.70 ng/g. By contrast, *Percent Time in Mixed-Sex Groups* and *Space Experience Outdoors at Night* increase FGMs: a 10% increase in time produces a 1.90 ng/g increase, and a 5000 ft^2^ increase in space experience produces a 0.60 ng/g in FGMs.

Table 8 presents univariate regressions of the independent variables and the CV of FGMs. Associated with lower variability of FGMs were *Enrichment Diversity, Social Group Contact* and *Social Experience, Percent Time with Juveniles,* both *Space Experience at Night* variables, *Mean Daily Walking Distance, Feeding Predictability* and *Latitude of Zoo.* The variable associated with increased variability was *Percent Time with In/Out Choice.*

**Table 8.** Univariate linear regressions between CV of fecal glucocorticoid metabolite concentrations and previously published risk factors (independent variables) for Asian and African elephants in North American zoos that participated in the Elephant Welfare Project. Variables entered into multi-variable analyses (P<0.15) are **bolded**.

| **Independent variable** | **N** | **Beta** | **SE** | **P value** |
| --- | --- | --- | --- | --- |
| Percent Time in Mixed Sex Herds (unpublished) | 237 | -0.015 | 0.022 | 0.507 |
| **Enrichment Diversity** | **222** | **-14.524** | **4.566** | **0.002** |
| Alternate Feeding Methods | 231 | -2.216 | 3.421 | 0.518 |
| **Social Group Contact** | **237** | **-0.451** | **0.135** | **0.001** |
| Walking (14 or more hours per week) | 217 | -0.342 | 0.470 | 0.468 |
| Feeding Diversity | 224 | -1.395 | 3.008 | 0.643 |
| Sex (ref: male) | 237 | -0.790 | 1.894 | 0.677 |
| Percent Time Managed | 218 | 0.022 | 0.040 | 0.580 |
| **Percent Time with Juveniles** | **237** | **-0.042** | **0.021** | **0.044** |
| Percent Time Housed Separately | 237 | -0.004 | 0.022 | 0.858 |
| Transfers | 237 | 0.411 | 0.363 | 0.260 |
| **Percent Time In/Out Choice** | **237** | **0.102** | **0.035** | **0.004** |
| **Social Experience** | **237** | **-0.830** | **0.370** | **0.026** |
| Recumbence Rate | 63 | 0.229 | 0.465 | 0.625 |
| > 0 Percent Time on Hard Substrate | 237 | -0.012 | 0.060 | 0.838 |
| > 0 Percent Time Soft Substrate | 237 | 0.039 | 0.056 | 0.486 |
| **Space Experience Outdoors Night** | **237** | **-0.016** | **0.009** | **0.076** |
| Space Experience In/Out Choice (per 500 ft<sup>2</sup>) | 237 | -0.016 | 0.015 | 0.304 |
| Joint Health: Absence or presence of joint abnormalities | 194 | 0.952 | 1.940 | 0.624 |
| **Space Experience Total Night (per 500 ft<sup>2</sup>)** | **237** | **-0.020** | **0.012** | **0.099** |
| Age of Elephant | 237 | 0.039 | 0.055 | 0.477 |
| **Mean Daily Walking Distance** | **60** | **-1.832** | **0.640** | **0.041** |
| **Feeding Predictability (ref: Unpredictable)** | **224** | **-2.564** | **1.145** | **0.026** |
| Mean Serum Cortisol | 215 | -0.023 | 0.116 | 0.844 |
| Keeper Attitude: Positive Opinions of Elephants | 192 | -1.561 | 1.641 | 0.343 |
| Keeper Attitude: Keeper as Herdmate | 192 | 1.373 | 1.356 | 0.312 |
| **Latitude of Zoo** | **237** | **-0.358** | **0.146** | **0.015** |
| Elephant Positive Behaviors | 160 | 1.363 | 1.010 | 0.179 |
| Elephant Interacts with Public | 160 | -0.822 | 1.106 | 0.458 |
| Species (ref=2, Asian) | 237 | -1.282 | 1.52 | 0.402 |

The multivariable model for CV of FGM (Table 9) indicates that *Percent Time In/Out Choice* increases FGM variability: when other variables are held constant, for each 10% increase in time there is a 0.9 % increase in CV of FGM. *Enrichment Diversity* and *Social Group Contact* both decreased variability. Each 1.0 increase in the Shannon Diversity Index of enrichment is associated with a 13.4% decrease in the CV of FGMs, and each additional *Social Group Contact* results in a 0.5% decrease. *Species* confounds *Enrichment Diversity* and *Social Group Contact* due to Asian elephants receiving, on average, slightly more enrichment than Africans (see Table 4), and Africans having contact with more social groups than Asians (Table 4), primarily because Africans are kept more often in larger groups.

**Table 9.** Multi-variable model of CV of fecal glucocorticoid metabolite concentrations for Asian (n=106) and African (n= 131) elephants in North American zoos that participated in the Elephant Welfare Project^1^.

| **Independent variable** | **Beta** | **SEM** | **P value** |
| --- | --- | --- | --- |
| Species<sup>1</sup> (ref: Asian) | 0.925 | 1.3855 | 0.504 |
| Sex (ref: female) | 0.828 | 1.7213 | 0.630 |
| Age | -0.050 | 0.0698 | 0.477 |
| Percent Time In/Out Choice | 0.090 | 0.0390 | 0.021 |
| Enrichment Diversity | -13.430 | 4.1904 | 0.001 |
| Social Group Contact | -0.516 | 0.0983 | 0.000 |

<sup>1</sup>Species is a confounder of *Social Group Contact* and *Enrichment Diversity*.

Because *Enrichment Diversity* is calculated on a zoo-level, Figure 1 shows the correlation between a zoo’s enrichment diversity score and the average CV FGM of the elephants at a zoo.

**Figure 1.**
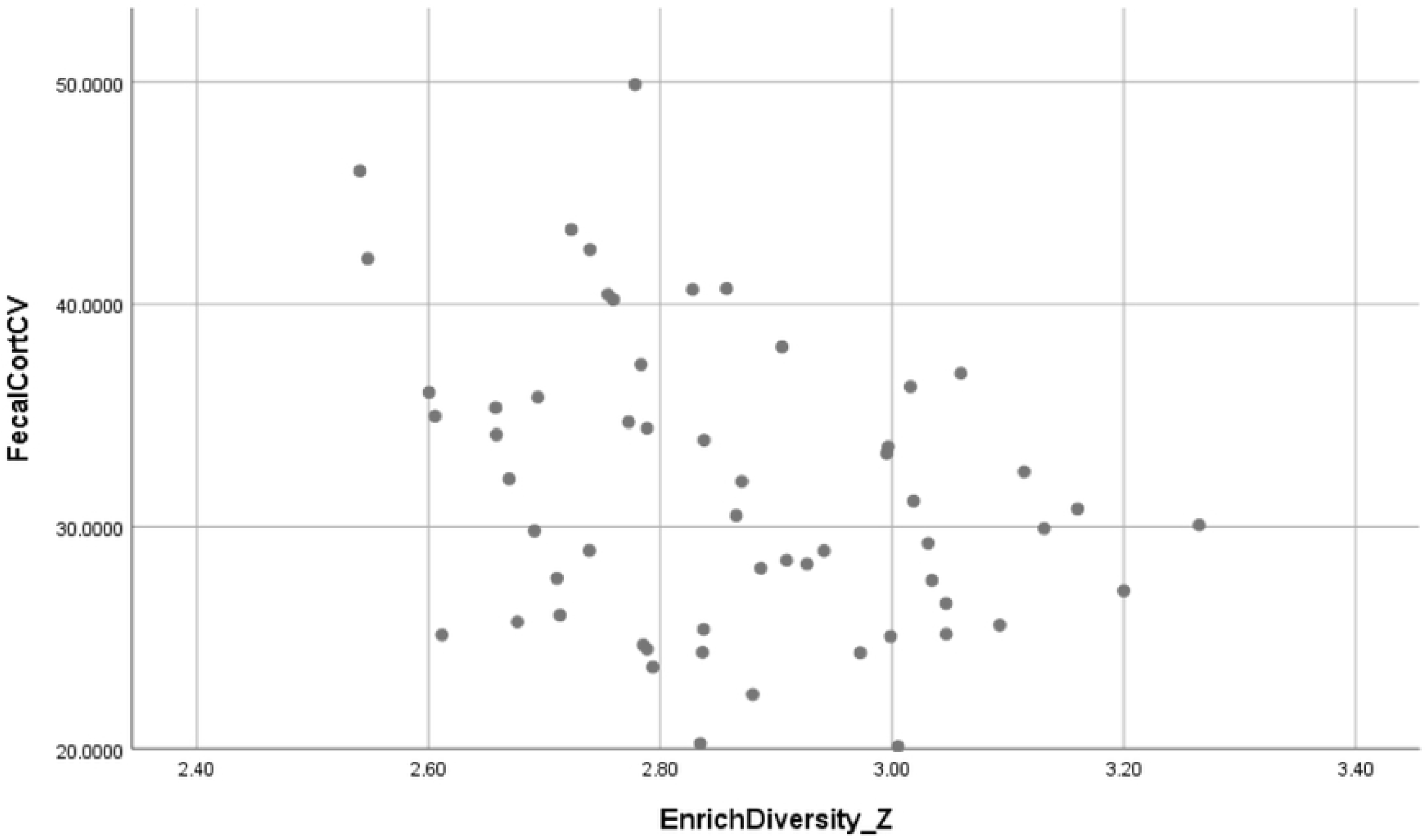
Correlation between zoos’ Enrichment Diversity scores and mean Coefficient of Variation of fecal corticoid metabolite concentrations at zoos (r = −0.339, n = 57, *P* = 0.010).

## Discussion

Epidemiological analyses of the EWP data point to a number of individual, social, housing and management factors that may affect adrenal activity in the zoo-housed population of elephants in North America. A higher risk of elevated FGM concentrations was found for Asian elephants with joint abnormalities, and African elephants housed in mixed-sex herds, whereas all elephants housed in northern latitudes had an increased risk of higher FGM in the multivariable models. More importantly, the results point to management factors that decrease FGMs in both species: having choice of being indoors and out, and management interactions with staff. The variability in FGM concentrations (CV) was reduced by enrichment and social groupings, and increased slightly by having a choice of indoor and outdoor spaces. Interestingly, walking distance and all three space experience variables were negatively correlated to FGM in Asian elephants, whereas for African elephants they were all positively associated. This pattern of correlations indicates that there are species differences in how housing space is experienced, which may suggest species-specific management protocols are needed, but could also be due to other covarying factors for each species.

Zoo elephants having the choice to be indoors or out appears to increase adrenal activity for both species, as indicated by significant negative associations between mean FGMs and the independent variables *Space Experience with In/Out Choice* (Asians) and *Percent Time with In/Out Choice (Africans).* The ability to actively move between spaces may stimulate the HPA axis in a positive way, or could be a source of stress if animals are moving to avoid negative states. Greco et al. [7] identified *Percent Time with In/Out Choice* as a risk factor for decreased frequency of nighttime stereotypy in the current population. Choice is generally beneficial to the welfare of captive animals because it increases an animal’s perceived control over its environment [42] and being given a choice of moving between indoor and outdoor areas at will has been associated with reduced stereotypic behaviors in polar bears [43], Asian elephants [44], and giant pandas [45]. However, in a separate epidemiological analysis of the current population [12], these same time and space choice variables were risk factors for increased foot and joint health problems, respectively. That ran counter to what was predicted. *Space Experience* and *Percent Time with In/Out Choice* are management variables that represent how much access an elephant has to indoor/outdoor areas, but not a measure of how much time an individual actually spends in either area or moving between them. Potentially an elephant with free access may choose to spend more time in smaller indoor areas (near keeper work areas) on hard substrate, thus contributing to foot and joint problems. Or it may be that elephants with a greater number of active pathologies are provided with more choice as a palliative treatment [12]. Powell and Vitale [44] reported that two of three Asian elephants given free access to indoor and outdoor areas at night preferred to be outdoors while the third individual stayed mostly indoors. Our results suggest that simply having choice may not always be stress-reducing and may depend on how much time an elephant actually spends in indoor and outdoor areas and under what circumstances.

Joint health was associated with FGM concentrations among Asian elephants. Elephants with joint problems had higher FGMs than those that did not, presumably due to pain. This was the case for zoo-housed Asian elephants, which spent more time on hard surfaces and were older on average than zoo-housed Africans [12], both risk factors for joint health problems, although there was no difference in the muscoskeletal scores assigned to individuals of these two populations [12]. This species difference may mean that joint health has been unintentionally diagnosed differently for each species, or is differentially experienced as more painful by Asian elephants.

*Latitude of Zoo* was a risk factor for higher FGMs in African elephants, increasing as a zoo location was more northwards. For Asians, this effect was only identified in the spring months. There are a variety of elephant management modifications that take place as seasons change, such as elephants spending more time confined inside or outside, with potential changes in social density or social contact that could account for increased social stress. Carlstead et al. [15] also found that *Latitude of Zoo* was a predictor of higher serum cortisol for the North American population of Asian elephants. Latitude as a risk factor of FGMs may be indicative of sensitivity to climatological changes that we would expect to be more pronounced the further north an elephant resides. Higher glucocorticoids have been reported during colder seasons among small numbers of zoo-housed Asian [46] and African [47] elephants. It remains unclear if latitude effects in the U.S. are due to climatological factors such as temperature and day length, or husbandry differences that cause more stressful conditions for elephants. In Thailand, mean FGM concentrations were ~28% higher in winter compared to the summer and rainy seasons, and were negatively associated with environmental factors: temperature and rainfall, but not humidity [48]. The need for more energy to maintain optimum body temperature and ensure survival in cooler temperatures could be related to this finding. This likely is an adaptive mechanism to ensure maintenance of anormal body temperature. In other ungulates, higher GC levels during winter have been found in white-tailed deer (*Odocoileus virginianus)* [49] and mule deer (*Odocoileus hemionus)* [50]. Elevated circulating GC levels as a response to cold stress also were documented in reindeer (*Rangifer tarandus)* [51] and in farm animals [52]. Seasonal trends in reproductive activity also have been documented. For example, a group of African elephants housed indoors because of cold temparatures at a zoo in Rhode Island showed prolonged non-luteal phases before re-initiating normal ovarian cycles in the spring [53] that could have been due to increased social stress or proximity effects, although GCs were not evaluated in that study.

There were three additional risk factors identified for African FGMs. First, *Percent Time Managed* by staff reduces FGMs, and also reduces daytime stereotypies for both species [7]. In Asians, there was a significant univariate correlation between FGMs and *Percent Time Managed,* but it did not make it into the multivariable model in this study. Therefore, stress in African elephants that is indicated by higher FGM concentrations and higher rates of stereotypy in the day time may be due to insufficient time spent in interactions with staff (i.e. cleaning and grooming, feeding, exercising and training). Positive interactions with keeper staff have been shown to be predictors of lower serum cortisol concentrations for both species [15]. The evidence points strongly to interactions with staff being stress-reducing, perhaps even calming for African elephants in zoos, and potentially for Asians as well.

Second, *Percent Time in Mixed-Sex Herds* is a small but identifiable factor in the lives of zoo-housed African elephants that is associated with increased FGMs. Social stress as measured by FGMs has been shown to be higher in free-ranging African elephant groups composed of non-related compared to related individuals [35], indicating that the composition of herds has effects on adrenal activity. It also should not be surprising to see elevated FGMs at institutions where there are bulls for breeding, a likely natural stressor. The third risk factor for African FGMs was *Space Experience Outdoors at Night*, which was associated with increased concentrations. There is no obvious explanation for why having more outdoor space at night would be associated with increased adrenal activity. Perhaps there are more social interactions occurring under the cover of darkness, without keepers nearby, which for some elephants might be stressful or, alternatively, stimulating. Posta et al. [54] reported that two zoo-housed African elephants spent a greater portion of their time outdoors at night walking, while others report significant social behaviors occurring during the night with free access to indoor and outdoor areas [55,56]. Holdgate et al. [8] also found that a subset of elephants from this population had a greater *Mean Walking Distance* if they had a greater *Space Experience at Night.* Therefore, evidence suggests that outdoor space at night facilitates activity of African elephants, and increased activity could account for the slight increase in FGMs identified in the multi-variable model.

The multi-variable model of CVs of FGMs revealed three risk factors; *Percent Time In/Out Choice, Enrichment Diversity* and *Social Group Contact.* Having more choice of being indoors or outdoors was associated with a slight increase in within-individual variability of FGMs, although the same variable was associated with reduced between-individual mean FGMs for African elephants. Therefore, while the overall population effect of choice appears to be stress-reducing, it leads to slightly increased variability in individuals. We speculate that this may be due to movements of other elephants in the herd going in and out in an unpredictable manner. A given individual might benefit from having increased choice and control over its own situation, but it has no control over the whereabouts of other elephants, potentially resulting in more variable stress responses. Cochrem [57] points out that CV needs to be included in studies of GCs because the factors that account for within-individual variation and their adaptive significance for a species, such as personality, coping styles, genetic or maternal influences, are little known for most species. For example, increased variability in FGMs was correlated with abnormal reproductive function, rates of fighting, and institutional mortality rates in rhinoceros [39], leading to the conclusion that the variability of FGMs is a valuable measure of stress responsiveness that has biological costs to the animal. The subject of individual variation in GC responses to stressors has included studies investigating different coping styles and disease susceptibility [58], and a better understanding of inter- and intra-individual variation in HPA reactivity would be beneficial to our use of GCs as a welfare measure [16].

*Enrichment Diversity* was strongly associated with a reduction in CV of FGMs, but not with mean FGMs, suggesting that having multiple enrichment options functions to moderate adrenal reactivity of individuals. Brown et al. [5] found enrichment diversity to be positively correlated with reproductive health in African females of the EWP, both in terms of reduced acyclicity and normalization of prolactin secretion, and our results suggest that diverse enrichment is an important management factor for zoo elephant welfare. Although enrichment has been shown to reduce GCs in rhesus monkeys [59] or rodents [i.e. 60], such demonstrations compare animals with and without enrichment under slightly stressful caging situations, demonstrating that enrichment facilitates coping with stress. However, all elephants of the EWP received some form of enrichment at their zoo, and the frequency with which different enrichments were provided was found to impact the variability of FGMs within but not between individuals. In an experimental study of mice that provided three different levels of enriched housing, mice housed in stress-reduction, “calm” cages consisting of a large cage with a cardboard nest box, paper nesting material, and a tube, exhibited significant and lasting reductions over time in FGM levels compared to mice housed in less enriched, standard caging [61]. Hence, ideally-enriched caging produced permanent calming effects on mice. In our analysis, *Enrichment Diversity* scores of zoos were derived from surveys of zoo managers providing the percentage of days their elephants had access to 30 different types of enrichment items, ranging from exhibit features such as sand or dirt piles, mud wallows, pools, logs, scratching posts and sprinklers, to the provision of manipulatable objects such as balls, tires and hanging objects, to feeding items such as browse and treat boxes/bags, and scents, music and problem-solving tasks [6]. We found the zoo average FGM CVs to be negatively correlated with the frequency of only three of the 30 enrichment types: problem-solving (r = −0.348, n = 57, p = 0.007), hanging objects (r = −0.261, p = 0.048) and scratching posts (r = −0.340. p = 0.009); three enrichments that intensely engage elephants. All evidence together strongly suggests that enrichment has a “calming” effect on stress responses of elephants, most likely by providing additional behavioral options and/or cognitive opportunities to cope with their daily lives.

Last, being a member of more social groups (*Social Group Contact)* also was associated with lower variability in FGMs. Therefore, being a familiar and accepted member of multiple social groups may also stabilize HPA-axis activity in a manner similar to *Enrichment Diversity*, effectively increasing social enrichment diversity, a clear benefit for elephant welfare.

## Conclusions

Results elucidate species differences in adrenal responses of elephants in zoo environments. African elephants appear to be more responsive to social stressors than Asians. It is well known that Asian elephants are not as bonded to large social groups as their African cousins and, therefore, have more limited hierarchical stratification, whereas African elephants live and interact in multi-tiered groups with presumably more social constraints [62]. Another species difference is that Asians might be more sensitive to stress caused by joint pain than Africans, but rates of joint problems and age differences between the two populations complicate this conclusion. In any case, the evidence points to poor joint health being a stress-related welfare problem for the U.S. population of Asian elephants. For both species, zoos located at more northern latitudes were characterized by elephants with slightly to significantly higher FGMs. It is unclear if these responses are due to climatological or management factors, or both. One factor that reduced FGMs for both species was more time being managed, suggesting time spent with keepers has a positive effect. More time being managed also was associated with reduced stereotypy [7], perhaps related to less boredom. Finally having diverse enrichment options and contact with multiple social groups also appears to be calming for elephants, reducing intra-individual variability in FGMs. Together, all evidence points to the beneficial effects of diverse enrichment opportunities, including cognitive enrichment for zoo-housed elephants. We conclude that there are many avenues for further research on stress in zoo-housed elephants, and monitoring FGMs longitudinally is a proven non-invasive method for determining factors contributing to adrenal function, stress and coping responses in elephants. The species differences observed in adrenal responses to zoo factors suggests that a one-size-fits-all management strategy may not be the best for both Asian and African elephans, and that more species-specific approach to husbandry may be needed.

## Acknowledgments

This work was part of a large-scale project “Using Science to Understand Zoo Elephant Welfare”, funded by a 2010 National Leadership Grant from the Institute for Museum and Library Services (IMLS) (grant no. LG-25-10-0033-10) from the U.S. IMLS provided financial support only for the conduct of the research and preparation of this article. The authors would like to acknowledge the significant efforts of the IMLS study principal investigators (Kathy Carlstead, Janine Brown, Nadja Wielebnowski, David Shepherdson, Mike Keele, Anne Baker, Beth Posta, Joy Mench, Candice Dorsey). In addition, we thank Steve Paris for conducting the FGM analyses. Special thanks to the project manager Cheryl Meehan and statistics advisor Jen Hogan. We are grateful to the AZA Elephant TAG and Vistalogic, Inc. for technical support and software services. Finally, sincere thanks to the people and elephants at each of the following zoos for incredible participation and support of the project: African Safari, Albuquerque Biological Park, Audubon Institute, Birmingham Zoo, BREC’s Baton Rouge Zoo, Buffalo Zoological Gardens, Busch Gardens, Buttonwood Park Zoo, Caldwell Zoo, Calgary Zoo, Cameron Park Zoo, Cheyenne Mountain Zoological Park, Cincinnati Zoo & Botanical Garden, Cleveland Metroparks Zoo, Columbus Zoo, Dallas Zoo, Denver Zoo, Dickerson Park Zoo, Disney’s Animal Kingdom, El Paso Zoo, Fresno Chaffee Zoo, Greenville Zoo, Honolulu Zoo, Houston Zoological Gardens, Indianapolis Zoological Society, Inc., Jacksonville Zoological Gardens, Knoxville Zoological Gardens, Lee Richardson Zoo, Little Rock Zoological Garden, Los Angeles Zoo and Botanical Gardens, Louisville Zoological Garden, Lowry Park Zoological Garden, Maryland Zoo, Memphis Zoological Garden and Aquarium, Metropolitan Toronto Zoo, Milwaukee County Zoological Gardens, Montgomery Zoo, Nashville Zoo, National Zoo, Niabi Zoo, North Carolina Zoological Park, Oakland Zoo, Oklahoma City Zoological Park, Oregon Zoo, Parque Zoologico de Leon, Phoenix Zoo, Point Defiance Zoo and Aquarium, Reid Park Zoo, Riverbanks Zoological Park, Roger Williams Park Zoo, Rosamond Gifford Zoo at Burnet Park, San Antonio Zoological Gardens & Aquarium, San Diego Safari Park, San Diego Zoo, Santa Barbara Zoological Gardens, Sedgwick County Zoo, Seneca Park Zoo, Saint Louis Zoo, The Kansas City Zoo, Toledo Zoo, Topeka Zoological Park, Tulsa Zoological Park, Utah’s Hogle Zoo, Virginia Zoological Park, Wildlife Conservation Society—Bronx Zoo, Wildlife Safari, Woodland Park Zoo, Zoo Atlanta, Zoo de Granby, Zoo Miami.

